# Neuronal firing rates diverge during REM and homogenize during non-REM

**DOI:** 10.1101/069237

**Authors:** Hiroyuki Miyawaki, Brendon Watson, Kamran Diba

## Abstract

Neurons fire at highly variable innate rates and recent evidence suggests that low and high firing rate neurons display different plasticity and dynamics. Furthermore, recent publications imply possibly differing rate-dependent effects in hippocampus versus neocortex, but those analyses were carried out separately and with possibly important differences. To more effectively synthesize these questions, we analyzed the firing rate dynamics of populations of neurons in both hippocampal CA1 and frontal cortex under one framework that avoids pitfalls of previous analyses and accounts for regression-to-the-mean. We observed remarkably consistent effects across these regions. While rapid eye movement (REM) sleep was marked by decreased hippocampal firing and increased neocortical firing, in both regions firing rates distributions widened during REM due to differential changes in high-firing versus low-firing cells in parallel with increased interneuron activity. In contrast, upon non-REM (NREM) sleep, firing rate distributions narrowed while interneuron firing decreased. Interestingly, hippocampal interneuron activity closely followed the patterns observed in neocortical principal cells rather than the hippocampal principal cells, suggestive of long-range interactions. Following these undulations in variance, the net effect of sleep was a decrease in firing rates. These decreases were greater in lower-firing hippocampal neurons but higher-firing frontal cortical neurons, suggestive of greater plasticity in these cell groups. Our results across two different regions and with statistical corrections indicate that the hippocampus and neocortex show a mixture of differences and similarities as they cycle between sleep states with a unifying characteristic of homogenization of firing during NREM and diversification during REM.

**Significance Statement:** Miyawaki and colleagues analyze firing patterns across low-firing and high-firing neurons in the hippocampus and the frontal cortex throughout sleep in a framework that accounts for regression-to-the-mean. They find that in both regions REM sleep activity is relatively dominated by high-firing neurons and increased inhibition, resulting in a wider distribution of firing rates. On the other hand, NREM sleep produces lower inhibition, and results in a more homogenous distribution of firing rates. Integration of these changes across sleep results in net decrease of firing rates with largest drops in low-firing hippocampal pyramidal neurons and high-firing neocortical principal neurons. These findings provide insights into the effects and functions of different sleep stages on cortical neurons.

## Introduction

Firing rates vary among neurons and across time. The dynamic range of a neuron’s firing is determined by a combination of membrane geometry, distribution and subtypes of ion channels, and synaptic efficacy (Koulakov et al., 2009; Yassin et al., 2010; Roxin et al., 2011; Lim et al., 2015; Stuart and Spruston, 2015; Nigam et al., 2016). Changes in these properties can potentially alter a neuron’s gain function or “excitability,” altering the neuron’s encoding properties (Cheng and Frank, 2008; Lee et al., 2012). Recent evidence suggests that a neuron’s firing rate is also homeostatically regulated (Vyazovskiy et al., 2009; Hengen et al., 2013; Miyawaki and Diba, 2016; Watson et al., 2016), and that modifications in membranes and synapses can work to maintain the neuron’s dynamic range (Marder and Goaillard, 2006; Turrigiano, 2011). Several studies from different labs indicate that these modifications are at least partially state-dependent; the emerging picture is that firing rates of neurons increase during waking (Vyazovskiy et al., 2009; Hengen et al., 2016; Miyawaki and Diba, 2016; Watson et al., 2016) and decrease during sleep (Vyazovskiy et al., 2009; Grosmark et al., 2012; Miyawaki and Diba, 2016; Watson et al., 2016), in a perpetual dance around a dynamic range.

The various waking and sleep states feature different activity levels of the neuromodulatory systems, which contribute uniquely to the excitability of neuronal circuits, network firing patterns, and the plasticity of their synapses (Hobson and Pace-Schott, 2002; Brown et al., 2012). For example, REM is characterized by high acetylcholine and low noradrenaline, serotonin and histamine levels, while waking and NREM respectively feature high and low levels of these neuromodulators (Hobson and Pace-Schott, 2002; Brown et al., 2012). Unique brainstem and thalamocortical networks are also active within each state, producing state-specific oscillatory firing patterns (Saper et al., 2010; Brown et al., 2012; Weber and Dan, 2016). The differing neuromodulatory and network backgrounds lead to different overall firing rates in REM, NREM, and waking (Vyazovskiy et al., 2009; Miyawaki and Diba, 2016; Watson et al., 2016), but averaging can also mask significant variations within each state (Grosmark et al., 2012; Miyawaki and Diba, 2016; Watson et al., 2016).

It was recently shown that sleep yields a net decrease in the firing rates of both hippocampal (Miyawaki and Diba, 2016) and frontal cortex neurons (Watson et al., 2016). These changes were likely explained by synaptic downscaling (Tononi and Cirelli, 2014), triggered in the hippocampus by sharp-wave ripples and sleep spindles during NREM sleep, and incorporated over the course of REM sleep (Miyawaki and Diba, 2016) and in the neocortex, triggered by alternating cycles of UP/DOWN states (Bartram et al., 2017; Gulati et al., 2017). In Miyawaki and Diba (2016) and Watson et al. (2016), we took trouble to evaluate firing rate changes between different epochs of the same state (e.g. NREM_i_ and NREM_i+1_ epochs in sleep) to avoid confounds of state-dependent neuromodulation. However, some questions remain regarding how firing patterns of neurons of differing excitabilities change within each of these states and on transitions between these states, and how these compare between hippocampal and neocortical neurons. In particular, low and high firing neurons, with presumed low and high levels of excitability, are expected to be affected differently by activity-driven homeostasis and bear differing levels of plasticity (Koulakov et al., 2009; Hengen et al., 2013; Lim et al., 2015; Grosmark and Buzsaki, 2016). While this question was addressed to some extent in our previous work, understanding such effects is complicated by regression to the mean (RTM), for which the null hypothesis allows that firing rates of low-firing neurons should increase and those of high-firing neurons should decrease across any two comparative periods. In this report, we investigate changes in firing rates of neurons within different stages of sleep and the effects of transitions between sleep stages in both hippocampus and frontal cortex, while carefully controlling for RTM. We find that transitions to REM and NREM sleep states differentially affect low-firing and high-firing neurons in each state. In both hippocampus and frontal cortex, we find that REM sleep is marked by inhibition, and the spread between low- and high-firing neurons increases, while NREM results in a more homogenized and narrowed distribution of rates. These observations may help provide insights into the function and effects of sleep states on cortical networks of neurons.

## Materials and Methods

We re-analyzed data previously recorded from hippocampal CA1 region of four male rats (Miyawaki and Diba, 2016; Miyawaki et al., 2017) and frontal cortex of 11 male rats (Watson et al., 2016). Details of the experimental protocols, including animals, surgery, electrophysiological recoding, spike detection and clustering, and sleep detection can be found in these refs (Miyawaki and Diba, 2016; Watson et al., 2016; Miyawaki et al., 2017) and are summarized below. EMG was obtained from either the nuchal muscles or from correlated high-frequency (300 - 600 Hz) signals from brain electrodes (Schomburg et al., 2014; Watson et al., 2016). All experimental procedures were in accordance with the National Institutes of Health guidelines and approved by the University of Wisconsin-Milwaukee, New York University, and Weill Cornell Medical College Institutional Animal Care and Use Committees. Numbers of analyzed cells and states/transitions for each dataset are summarized in Table 1. To estimate firing rates reliably, only NREM > 150 s and REM > 100 s were used for all analyses.

**Table 1.** Number of cells and states/transitions. Numbers for time normalized analyses (Fig. 1, 2 and 6A,B) and for *CI*/*DI* analyses (Fig. 4,5, and 6C-F) are shown in top and bottom of each cell.

| States/transition type | Hippocampus |  | Frontal cortex |  |
| --- | --- | --- | --- | --- |
|  | Pyramidal cells | Interneurons | Principal cells | Interneurons |
| NREM | 25717 cell-epochs in 462 epochs | 3049 cell-epochs in 406 epochs | 8097 cell-epochs in 232 epochs | 785 cell-epochs in 205 epochs |
|  | 25466 cell-epochs in 458 epochs | 3045 cell-epochs in 405 epochs | 7977 cell-epochs in 230 epochs | 735 cell-epochs in 203 epochs |
| REM | 15158 cell-epochs in 277 epochs | 1783 cell-epochs in 241 epochs | 3999 cell-epochs in 123 epochs | 373 cell-epochs in 109 epochs |
|  | 13106 cell-epochs in 244 epochs | 1615 cell-epochs in 221 epochs | 3463 cell-epochs in 109 epochs | 296 cell-epochs in 96 epochs |
| **NREM-REM** | 13159 cell-epochs in<br>240 transitions<br>11312 cell-epochs in<br>208 transitions | 1528 cell-epochs in<br>210 transitions<br>1334 cell-epochs in<br>188 transitions | 3380 cell-epochs in<br>105 transitions<br>2575 cell-epochs in<br>85 transitions | 319 cell-epochs in<br>93 transitions<br>222 cell-epochs in<br>76 transitions |
| **REM-NREM** | 10355 cell-epochs in<br>190 transitions<br>8105 cell-epochs in<br>155 transitions | 1237 cell-epochs in<br>168 transitions<br>992 cell-epochs in<br>138 transitions | 3493 cell-epochs in<br>103 transitions<br>2995 cell-epochs in<br>87 transitions | 323 cell-epochs in<br>92 transitions<br>255 cell-epochs in<br>77 transitions |
| **Wake-NREM** | NA<br>4176 cell-epochs in<br>80 transitions | NA<br>724 cell-epochs in<br>100 transitions | NA<br>1697 cell-epochs in<br>48 transitions | NA<br>194 cell-epochs in<br>43 transitions |
| **NREM-WAKE** | NA<br>1014 cell-epochs in<br>29 transitions | NA<br>252 cell-epochs in<br>37 transitions | NA<br>408 cell-epochs in<br>28 transitions | NA<br>29 cell-epochs in<br>transitions |
| **REM-WAKE** | NA<br>298 cell-epochs in<br>20 transitions | NA<br>89 cell-epochs in 8<br>transitions | NA<br>60 cell-epochs in 28<br>transitions | NA<br>2 cell-epochs in 2<br>transitions |
| **NREM-REM-**<br>**NREM** | 8767 cell-epochs in<br>165 triplets<br>6769 cell-epochs in<br>133 triplets | 1017 cell-epochs in<br>145 triplets<br>776 cell-epochs in<br>116 triplets | 2926 cell-epochs in<br>88 triplets<br>2123 cell-epochs in<br>66 triplets | 271 cell-epochs in<br>78 triplets<br>173 cell-epochs in<br>58 triplets |
| **REM-NREM-**<br>**REM** | 5031 cell-epochs in<br>92 triplets<br>3257 cell-epochs in<br>60 triplets | 609 cell-epochs in<br>79 triplets<br>454 cell-epochs in<br>54 triplets | 1098 cell-epochs in<br>35 triplets<br>443 cell-epochs in<br>19 triplets | 98 cell-epochs in 3<br>triplets<br>41 cell-epochs in 1<br>triplets |
| **SLEEP** | NA<br>4983 cell-epochs in<br>88 sleeps | NA<br>585 cell-epochs in<br>74 sleeps | NA<br>1379 cell-epochs in<br>46 sleeps | NA<br>129 cell-epochs in<br>35 sleeps |
| **WAKE-**<br>**SLEEP-WAKE** | NA<br>648 cell-epochs in<br>24 sequences | NA<br>304 cell-epochs in<br>37 sequences | NA<br>758 cell-epochs in<br>35 sequences | NA<br>71 cell-epochs in 2<br>sequences |

### Time normalized mean firing rates

In one series of analyses of sleep sequences (Fig 1B,E; Fig 2B,E; Fig 6A,B) NREM and REM epochs were divided into 30 bins and 10 bins, respectively, since NREM epochs are generally longer. Additionally, cells were sorted into quintiles within each epoch and firing rates of cells in each quintile were calculated in each bin. Importantly, sorting was based on their mean firing rates over the entirety of the windows of interests depicted in each panel, to avoid RTM effects which are particularly evident when tracking ranked groups of units such as quintiles. While ranking was based on the entire epoch in these analyses, to allow for comparisons across quintiles, firing rates were normalized by the mean firing rate in the last one-third of the first state.

**Figure 1.**
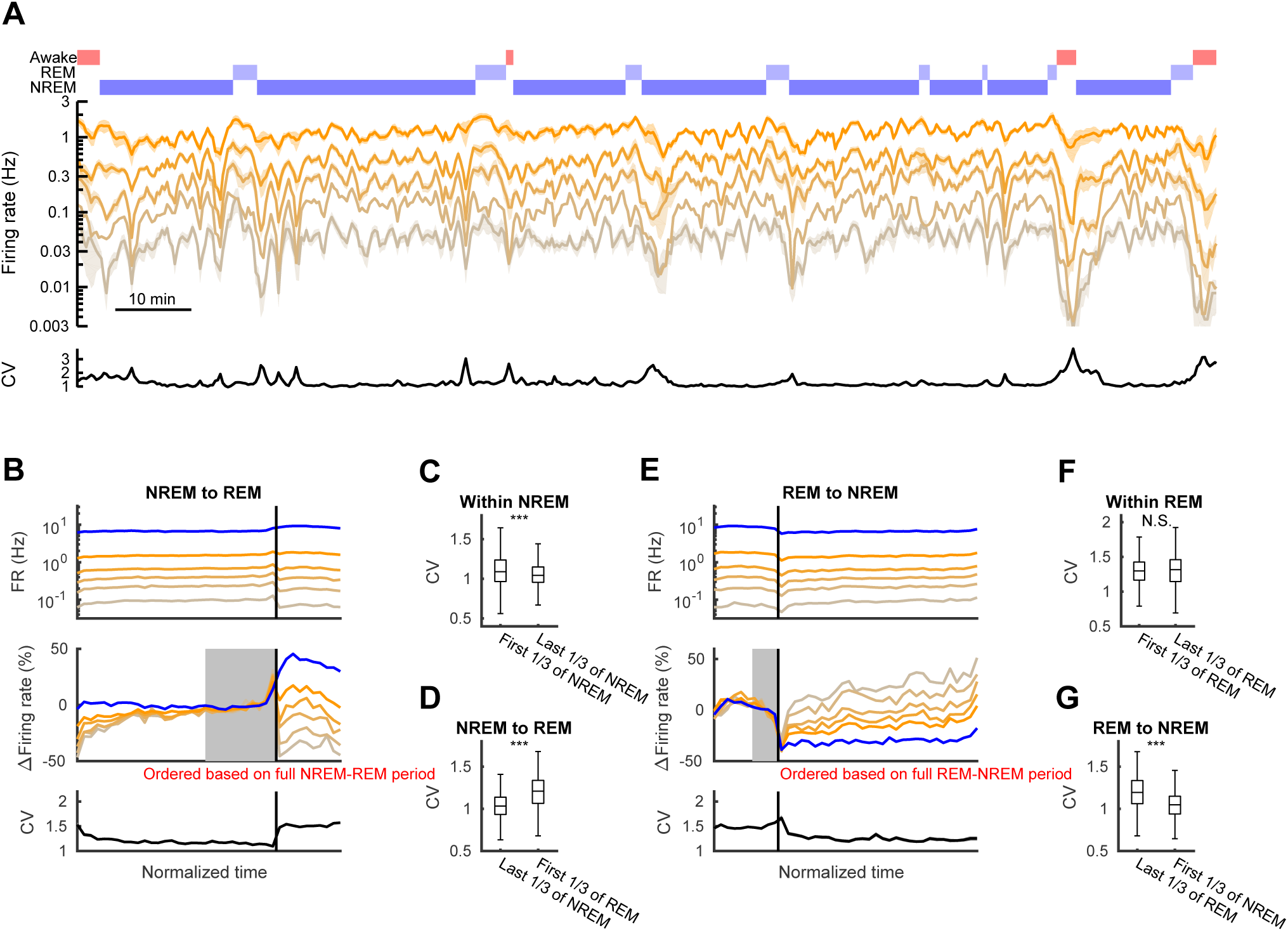
Firing rates of hippocampal neurons over transitions between sleep states. (A) An example period showing mean firing rates of hippocampal pyramidal cells (n=94) sorted into five quintiles in sliding 1-min windows (20 s steps). The hypnogram and coefficient of variation (CV) are shown at top and the bottom, respectively. (B) Mean firing rates of each quintile of pyramidal cells (yellow lines) and interneurons (blue line) over transitions (vertical black line) from NREM to REM pooled across recordings (top panel). Quintiles were sorted independently for each analyzed NREM-REM doublet prior to averaging for presentation (see Table 1 for details). The middle panel shows firing rates normalized to mean across full period shown and represent the relative change from the mean in the last third of the NREM epoch (the period indicated in gray). The bottom panel shows the mean CV of the complete distribution of pyramidal cells. (C, D) CV changes within NREM and on the transitions from NREM to REM. Significance was based on the Wilcoxon rank sum test. Error bars and line shades indicate SEM. *** p<0.001, N.S., not significant. (E-G) Same with (B-D), but for transitions from REM to NREM. Alignment in E based on last third of REM (gray).

**Figure 2.**
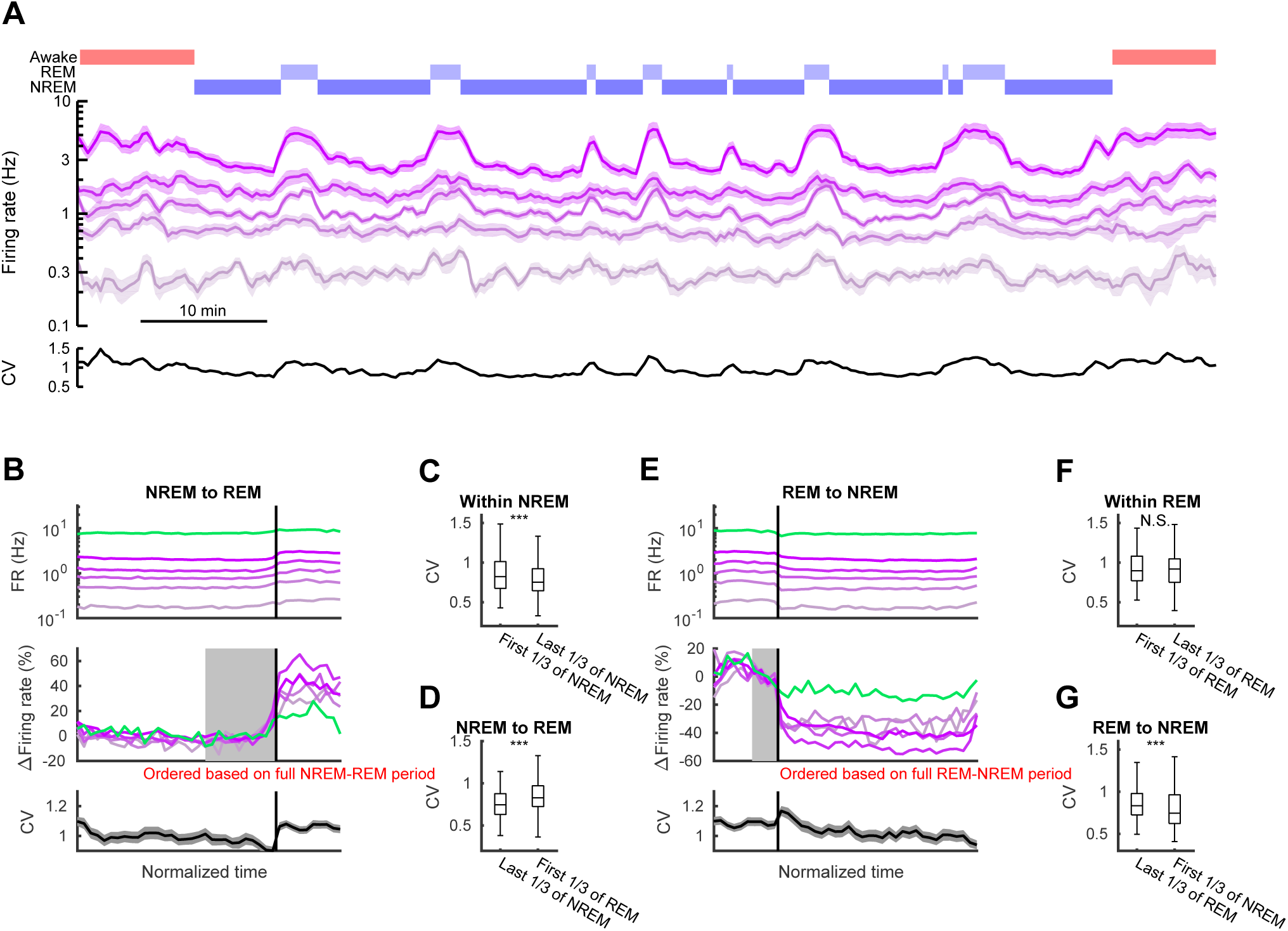
Firing rates of frontal neocortical neurons over transitions between sleep states. Same as in Figure 1, but for frontal cortex. (A) An example period showing firing rates of frontal neocortical principal neurons (n=89) in five quintiles (sliding 1-min windows with 20 s steps). Hypnogram and coefficient of variation (CV) shown at top and the bottom, respectively. (B, E) Firing rates of principal neurons (purple; top panels) and interneurons (green) over NREM-REM (B) and REM-NREM (E) transitions. Firing rates in the middle panels were normalized to the mean across full period shown and aligned to mean in the last third of NREM in (B) and last third of REM in (E) (gray regions). CV of principal neuron firings on bottom panels. (C, D, F, G) CV changes within states and across sleep state transitions. Significance was based on the Wilcoxon rank sum test. Error bars and line shades indicate SEM. *** p<0.001, N.S., not significant.

### Change index and deflection index

For a second set of analyses, change index (*CI*) for a quintile was defined as 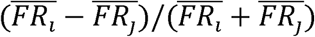, where *FR*_*i*_ and *FR*_*j*_ are mean firing rates of a neuron over time periods *i* and *j* (*i < j*), respectively, and 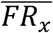 is the mean of the mean *FR*_*x*_ over the neurons in the quintile. Neurons were first separated into quintiles based on *FR*_*i*_ within each epoch, and quintile *CI* was then calculated across a given pair of epochs. If more than 20% of cells did not fire in epoch *i* or *j*, that sequence was excluded from the analyses since *CI* of lowest quintile cannot be properly calculated in such a case. Note that this analysis still allows for changes in quintile membership between *i* and *j*. Because neuronal firing rates are log-normally distributed, the difference in logarithm of firing rates, Δ*log(FR)*, had been previously used to assess firing rate changes (Watson et al., 2016). Although *CI* and Δ*log(FR)* generally behave similarly, Δ*log(FR)* becomes singular when either *FR*_*i*_ or *FR*_*j*_ approach or equal zero. Therefore, in these analyses we opted to use *CI*.

Our null hypothesis was that changes across states are no different than changes within states, allowing for RTM. To evaluate the corresponding null distribution for this analysis, we generated 2000 shuffled surrogates by random flipping of *FR*_*i*_ and *FR*_*k*_ for each cell, where *k* is a control period of the same state as *i*. For analyses involving sleep (Fig 4; Fig 6C,D), these control periods were taken from the corresponding periods (e.g. first one-third; or entire epoch) in adjacent epochs of the same state (NREM or REM). Note that by chance these surrogate CIs therefore involve changes either forward or backward in time and always either ending or originating in epoch *i*. For transitions involving WAKE (Fig5; Fig 6F), control periods were randomly selected from 1-min periods of the same wake epoch (since wake periods separated by sleep display significantly different firing rates (Vyazovskiy et al., 2009; Miyawaki and Diba, 2016)). Shuffled mean and 95% confidence intervals of *CI* were obtained from this surrogate data. The deflection index (*DI*) was defined as difference of *CI* from the surrogate mean.

### Simulations

To better understand the behavior of *CI* and *DI* we generated three random datasets involving noise combined with no-change, additive firing rate increase, and multiplicative firing rate increase. Each dataset has 5000 cells and 3 epochs, corresponding to epochs *i*, *j*, and *k*. To mimic the variability of real data, first we set a baseline firing rate for each cell based on a log-normal distribution obtained from hippocampal pyramidal cells during NREM (mean = −0.4726 log_10_Hz, std= 0.4577 log_10_Hz) and then added random (“multiplicative”) noise proportional to each cell’s firing rate in each epoch (std=0.1403 log_10_Hz). For the no-change simulation, each cell kept the same baseline firing rates across epochs with only random noise producing fluctuation across epochs. For additive and multiplicative increase simulations, baseline firing rates in epoch *j* were increased (by addition of 0.05 Hz or multiplication by 1.1 for additive and multiplicative increases, respectively).

### Experimental Design and Statistical analyses

In this work we analyzed previously obtained data and no additional experiments were performed. Diversity of firing rates was evaluated by coefficient of variance (CV) and significance of difference was tested with Wilcoxon rank sum test. P-values of DIs were calculated relative to shuffled surrogates. Differences in DI and firing rate changes among quintiles were tested with one-way ANOVA. Firing rate distributions were compared by a Kolmogorov-Smirnov test. All analyses were performed with custom-written scripts running on MATLAB with statistics and machine learning tool boxes. Code is available upon request.

## Results

### Differential effects of REM and NREM on higher- and lower-firing rate hippocampal neurons

We previously recorded from populations of CA1 pyramidal cells and interneurons over multiple sleep and awake cycles (Miyawaki and Diba, 2016). Based on these data, we showed that mean firing rates in the hippocampal pyramidal cells increased within NREM but decreased through transitions between NREM and REM, and such zig-zag change resulted in net decrease across sleep (Grosmark et al., 2012; Miyawaki and Diba, 2016). However, it was not clear whether or not transitions between sleep states affect lower and higher firing cells uniformly. To address this question, we sorted the pyramidal cells into five quintiles based on their rank-ordered firing rates (Fig. 1A) and investigated their changes within and across NREM and REM sleep epochs (Fig. 1B-G). To overcome potential confounds from regression to the mean (RTM), here cells were sorted into quintiles according to their firing rates over the entirety of the periods shown in each panel (see Materials and Methods for further details). Although all quintiles showed gradual firing increases within NREM and sudden decreases at the transitions to REM, the relative magnitude of changes were different in lower- and higher-firing quintiles. Upon transitions to REM, lower firing cells showed large drops in activity, while higher firing cells showed little change (Δfiring rate = −41.5±3.3%, −31.0±2.8%, −14.6±2.9%, −5.9±2.5%, and 11.4±1.9% for low to high firing quintiles from last 1/3 of NREM to first 1/3 of REM, F_(4)_ = 59.2, p = 1.1×10^−45^, one-way ANOVA; Fig. 1B), effectively widening the distribution of firing rates and producing an increased coefficient of variation in the firing rates across neurons (ΔCV = 0.189±0.011, p = 1.0×10^−35^, Wilcoxon signed-rank test; Fig. 1B, D). Interneuron firing increased at the transition to REM (Δfiring rate = 38.8 ± 61.2%, p =1.1×10^−25^, Wilcoxon signed-rank test), consistent with a more competition-driven network (Xie et al., 2002). Within REM, firing rate changes were similar across quintiles and the CV did not change significantly (Fig. 1B,E,F). Upon the transition from REM to NREM, firing rates initially dropped in all quintiles. However, lower firing cells rebounded strongly (Fig. 1E) whereas higher firing cells were suppressed across NREM. Consequently, the CV of firing rates decreased (Fig. 1E,G, ΔCV = −0.155±0.014, p = 2.8×10^−21^, Wilcoxon signed-rank test), indicating a narrowed and more uniform distribution of neuronal firing rates. This rebalancing of excitability within NREM was accompanied by decreased firing in interneurons. In summary, we observed differential dynamics in lower and higher firing neurons, with transitions to REM widening the distribution of firing rates and both transitions to and continuation of NREM narrowing the distribution of firing rates.

### Differential effects of REM and NREM on higher- and lower-firing rate neocortical neurons

To examine whether these or similar state effects are also present in the neocortex, we extended these same analyses to neuronal spiking data recorded from frontal cortex of rats (Watson et al., 2016) and available on crcns.org (Fig. 2). Unlike in the hippocampus, firing rates increased at the transition from NREM to REM (Evarts, 1964; McCarley and Hobson, 1971; Vyazovskiy et al., 2009; Renouard et al., 2015; but also see Niethard et al., 2016). However, similar to the hippocampus, firing rate distributions widened upon this transition, with higher firing neocortical cells showing relatively larger increases at the transition to REM (Δfiring rate = 24.9±5.4%, 33.3±5.7%, 34.0±4.1%, 53.9±6.7%, and 41.3±3.8%, for each quintile from last 1/3 of NREM to first 1/3 of REM, F_(4)_ = 4.3, p = 0.002, one-way ANOVA; Fig. 2B), increasing the CV of firing rates (ΔCV = 0.033±0.021, p = 8.7×10^−5^, Wilcoxon signed-rank test; Fig. 2B,F) alongside increased firing in interneurons (Δfiring rate = 15.1 ± 48.7%, p =7.8 ×10^−4^, Wilcoxon signed-rank test). Upon transitions from REM to NREM, firing rates decreased in the neocortex, and as in the hippocampus (Fig. 2E,G), the decrease was stronger in higher-firing than in lower-firing neurons. There was an accompanied decrease in interneuron firing and a significant decrease in the CV of firing rates across neurons (ΔCV = −0.034±0.021, p = 5.9×10^−4^, Wilcoxon signed-rank test; Fig. 2E,G). The distribution of firing rates continued to narrow as NREM progressed (Fig. 2C). As the end of NREM approached, the firing rates began to increase, with further increases upon transition to REM. Overall, these results indicate that NREM and REM sleep states and transition between them have similar net effects on neurons with distributed firing rates in both the hippocampus and the neocortex.

### Regression-to-the-mean and sorting effects on firing rate changes

We wanted to better quantify these observations and to further evaluate how distributions of firing rates change within and across different sleep states. However, we first needed to better understand the relationship between variability and RTM in a population of neurons with log-normally distributed firing rates; ordering based on a part of analyzed data may bias the results due to RTM (Fig. 3A). We examined a simulated population of neurons where the source of variability is “multiplicative noise” proportional to each neuron’s firing rate (also see Materials and Methods) (Fig. 3B). Despite the absence of any change in our model population, lower-firing neurons show an apparent increased firing, while higher-firing neurons show an apparent decreased firing (Fig. 3C) when the quintiles were based on rank-ordering during the first epoch (i) of a sequence i-j. This is RTM and it can confound evaluations of true effects (e.g. from sleep). To control for RTM, we need to either rank-order cells by their mean firing rates over the entire sequence, as we did for analyses in Fig. 1-2, or else instate an appropriate correction. We introduced a shuffle correction in which we randomly flipped indices for epochs of the same state (e.g. *i* and *k* for a NREM_i_/REM_j_/NREM_k_) and repeated the analysis multiple times to obtain a surrogate distribution for the change index of quintiles (Miyawaki and Diba, 2016). This surrogate data provided us with valuable “control” shuffle means and confidence intervals for each quintile. We defined the “deflection index (*DI*)” as the difference between the observed change index (*CI*) and the surrogate mean within each quintile. These *DI*s were not significantly different from zero when changes were due only to noise (Fig. 3C).

**Figure 3.**
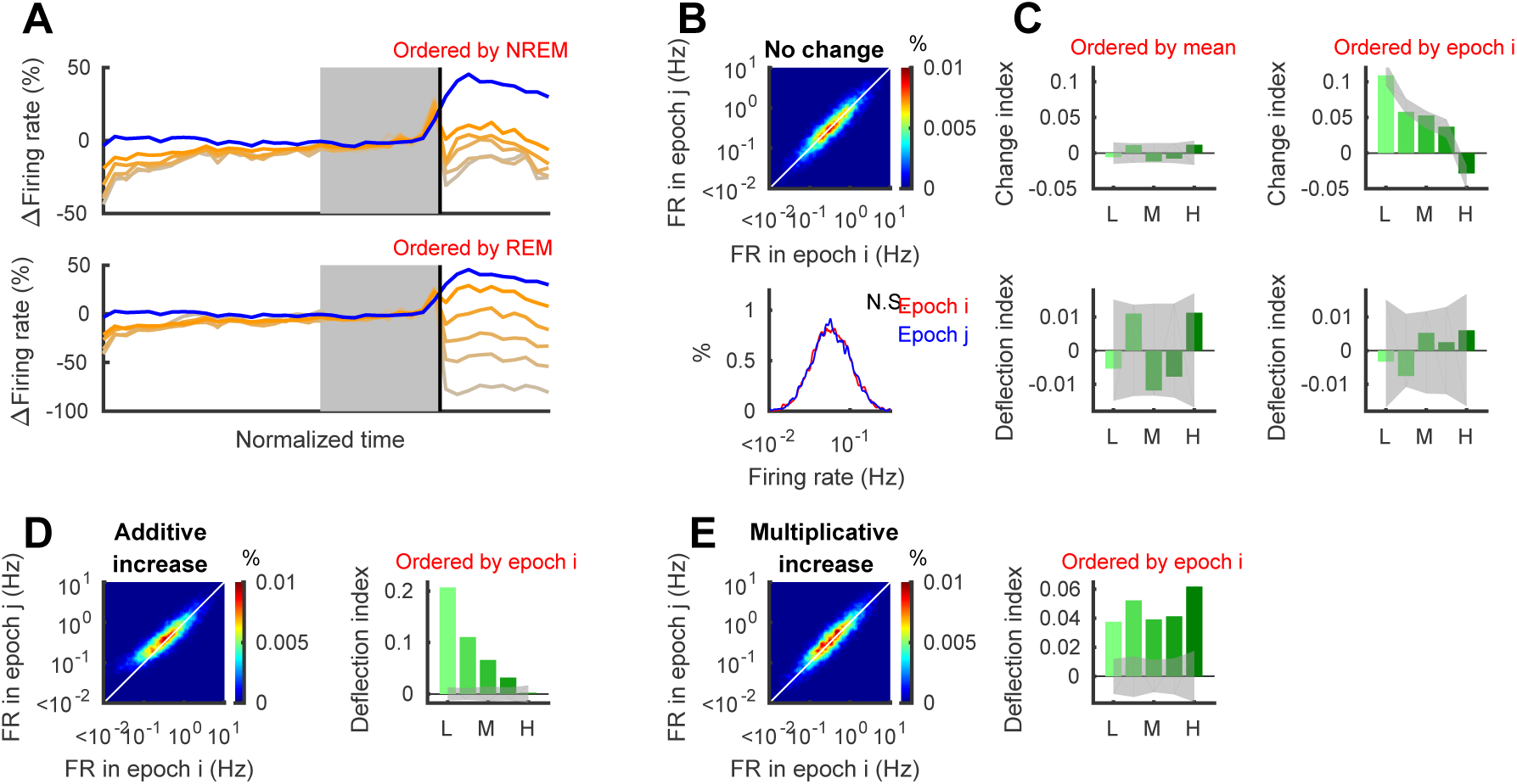
Deflection index can evaluate firing rate changes with correction of RTM. (A) Mean firing rates of hippocampal neurons in NREM-REM sequences as shown in figure 1B but different quintile separation (based on mean within NREM and REM for top and bottom, respectively). (B) Randomly generated firing rates with no change. (C) When cells ordered based on mean across states 1 and 2 (left column), there are no systematic difference on change index across quintiles. On the other hand, when cells ordered based on state 1 only (left column), systematic changes appear due to regression towards the mean. Deflection index (*DI*; for details see Material and Methods) can compensate the effect of RTM (bottom panels). (D,E) In case with changes in firing rate, *DI* is significantly different from zero. Here example of additive increase and multiplicative increase (E). Gray bands indicate 95% confidence intervals obtained from shuffling (2000 times). Each example has 5000 cells whose firing rates are distributed log-normally.

We then examined *DI*s under two scenarios with a simulated change in addition to noise: when firing rates increased across the population, either additively by a fixed amount for all cells (Scenario 1; Fig. 3D) or multiplicatively, by an amount proportional to each cell’s initial firing rate (Scenario 2; Fig. 3E). The shuffle-corrected *DI*s effectively described and differentiated the two scenarios. Under Scenario 1 the additive increase produced a larger relative effect on the *DI* in low-firing cells than in high-firing cells (Fig. 3C), while under Scenario 2, the evaluated *DI*’s correctly depicted a uniform increase across the population (Fig. 3E).

### Firing rate spread in REM and homogenization in NREM

We next used the RTM correction methods described above to enable comparisons of firing rates in pairs of epochs. We did these analyses either within each state of sleep or across different stages of sleep. This approach is complementary to that shown in Figures 1 and 2 and can serve as an independent verification of the observations shown there.

During NREM sleep, the average firing rates of hippocampal pyramidal neurons increased (mean *CI* = 0.058 ± 0.005, *p*=5.9×10^−27^, Wilcoxon signed-rank test). While the *CI* appeared to show the largest increase in low-firing cells, much of this apparent effect was due to RTM: in the shuffle corrected *DI*s, in fact the higher-firing quintiles showed the greatest relative increases (Fig. 4A, top row). In the frontal cortex the average *CI* decreased (mean *CI* = −0.023 ± 0.004, *p =*3.2×10^−9^, Wilcoxon signed-rank test). When cells were separated into quintiles, the *CI* appeared to indicate increased firing in lower-firing cells and decreased firing in higher-firing cells (Fig. 4A, bottom row), but those changes were also largely explained by RTM. After correction for RTM in the *DI*s, some increase was evident in the second lowest quintiles, along with a decrease in the two highest quintiles. Changes in firing rate distributions showed a consistent picture to the *DI* analyses (Fig. 4A, bottom) with the distribution for the hippocampus narrowing slightly and shifting towards increased firing, while that in the frontal cortex narrowing and shifting left towards decreased firing.

**Figure 4.**
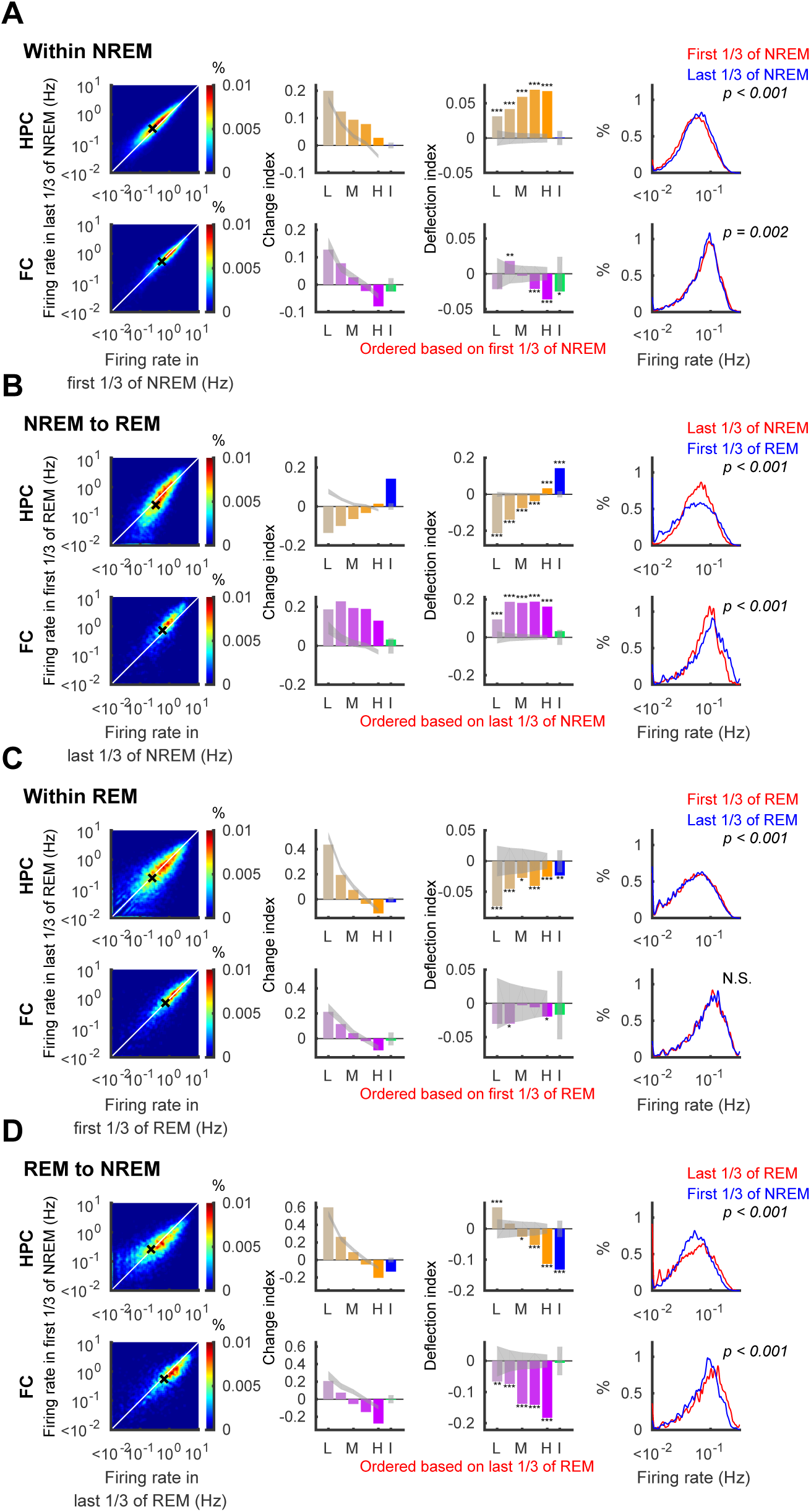
Firing rates diversify on transitions to REM and homogenize on transition to NREM. Firing rate changes within NREM (A), on transitions from NREM to REM (B), within REM (C), and on transitions from REM to NREM (D) in the hippocampus (HPC; top rows - orange) and the frontal cortex (FC; bottom rows - purple). Left panels show density (heat map) plots of firing rates. White lines indicate identity, and black crosses show means. Second and third panels illustrate change index (CI) and deflection index (DI) of each quintile of principal neurons (L: lowest quintile, M: middle quintile, H: highest quintile, yellow and purple bars) and interneurons (I, blue and green bars,) with 95% confidence interval (gray bands). Right panels show the firing rate distribution over all recorded principal neurons for the periods indicated in red and blue. P-values for the Kolmogorov-Smirnov test are indicated. * p<0.05, ** p<0.01, *** p<0.001.

At the transitions from NREM to REM, hippocampal pyramidal cells decreased firing in all quintiles but the highest one, with the largest decrease in the lowest quintile (*DI* = −0.213±0.019, −0.139 ± 0.016, −0076 ± 0.015, −0.038 ± 0.011, and 0.032 ± 0.010 for each quintile, all p values obtained by shuffling are <0.001; Fig. 4B, top row). Interneuron firing, on the other hand, increased (*CI* = 0.142 ± 0.013, p = 5.6×10^−22^, Wilcoxon signed-rank test), indicating a new steady state in the balance between network excitation and inhibition (but see Tsodyks et al., 1996). This increased inhibitory activity could potentially drive some of the decrease firing in pyramidal neurons (Niethard et al., 2016) and allow for a winner-take-all mechanism whereby some high-firing cells dominate REM dynamics at the expense of lower-firing cells. These dynamics were somewhat different for the frontal cortex, however; at the onset of REM, principal neurons in the frontal cortex increased firing across quintiles, while interneurons showed little change (*DI* = 0.094 ± 0.029, 0.187 ± 0.023, 0.181 ± 0.019, 0.188 ± 0.020 and 0.162 ± 0.016 for each quintile of principal neurons, p values < 0.001 relative to shuffles, *CI* = 0.031 ± 0.033 for interneurons, p=0.03, Wilcoxon signed-rank test). It is interesting to note however that the increased firing of hippocampal interneurons mirrored the overall increase in neocortical principal cell activity, consistent with neocortical control of hippocampal inhibition (Hahn et al., 2006; Basu et al., 2016). As a result of these changes, firing rate distributions became wider upon REM in both the hippocampus and the frontal cortex (Fig. 4B, right), consistent with our earlier analysis. Over the course of REM, we saw decreased firing across quintiles and interneurons in the hippocampus and in some quintiles of the neocortex (Fig. 4C). However, comparisons of the overall firing rate distributions did not reach statistical significance in the frontal cortex. The overall balance between excitation and inhibition therefore did not appear to change significantly within the course of REM states (Dehghani et al., 2016).

In contrast, when REM transitioned to NREM sleep, lower-firing quintiles showed increased firing while higher-firing quintiles and interneurons showed a firing decrease both in the hippocampus and in the frontal cortex (Fig. 4D). Interestingly, among the various dynamics we investigated only this transition from REM to NREM was marked by a renormalizing effect on firing rates across quintiles even after correction for RTM, and it was the only one we investigated that was marked by decreased firing of inhibitory cells in the hippocampus. NREM sleep therefore provided for the most uniform firing among the population of cells, potentially because of lower effective inhibition, whereas REM was marked by a widened distribution of firing activity.

### Neuronal firing changes at transitions to and from wake

Our results thus far have outlined the effects of transitions and continuation of REM and NREM sleep states on neurons at different levels of excitability. We next applied these same methods to analyze the effects of transitions between sleep and waking on different quintiles and focused on our corrected *DI* analysis. Immediately upon transitions from waking to NREM sleep (direct transitions from wake to REM are rarely observed), the hippocampus showed increases in the middle of the distribution (Fig. 5A) whereas the frontal cortex showed a decrease in high-firing cells. Nevertheless, the wake-to-sleep transition was accompanied by decreased inhibition in the hippocampus (*CI* = −0.076 ± 0.016, p=1.5×10^−6^, Wilcoxon signed-rank test) but not in the frontal cortex (−0.020 ± 0.032, p =0.60, Wilcoxon signed-rank test) and a narrowing of the distribution of firing rates in both regions (ΔCV = −0.708 ± 0.076, p = 2.4×10^−11^, and ΔCV = −0.239 ± 0.035, p = 6.1×10^−7^) for the hippocampus and the frontal cortex, respectively, Wilcoxon signed-rank test). The distribution narrowing in the hippocampus again indicates a new steady state in the balance between excitation and inhibition, with increased activity in the three middle quintiles (Fig. 5A), whereas the frontal cortex narrowing was a result of decreased firing in the highest-firing quintile and a trend towards more increase in progressively lower firing cells.

**Figure 5.**
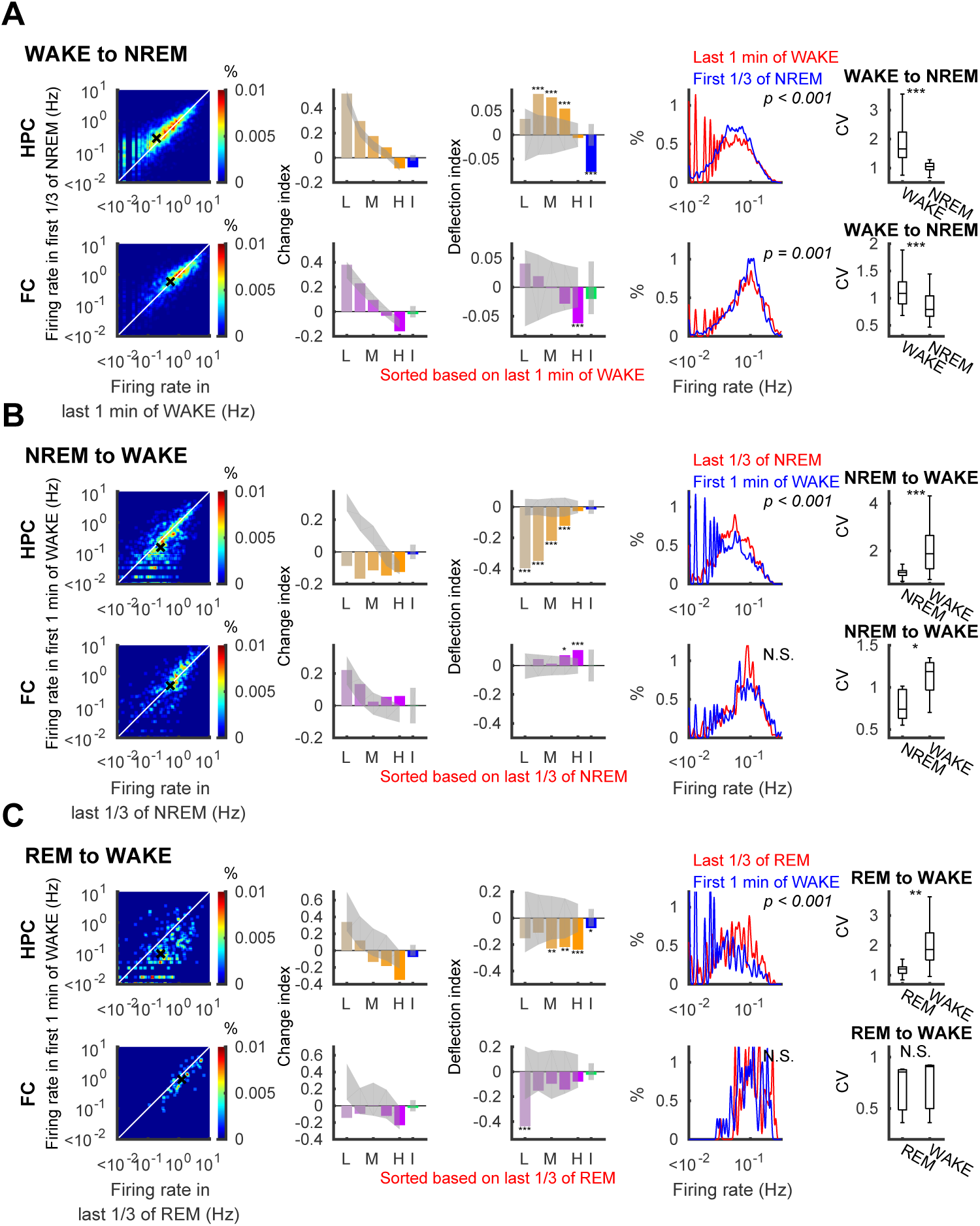
Firing rate changes at transitions between wake and sleep. Similar to Figure 4, firing rates (left panels), change (CI) and deflection (DI) indices (second and third panels) with 95% confidence intervals of shuffle mean (sheds on the bars), firing rate distribution (fourth panels) and coefficient of variation of firing rates (right panels) on transition from WAKE to NREM (A), NREM to WAKE (B), and REM to WAKE (C). Top and bottom rows in each panel present data from the hippocampus (HPC) and the frontal cortex (FC), respectively. P-values for Kolmogorov-Smirnov tests are indicated on the panels in the fourth column. L: lowest quintile, M: middle quintile, H: highest quintile, I: interneurons, * p<0.05, ** p<0.01, *** p<0.001.

In contrast, the distribution of firing rates widened at the onset of wake. At transitions from NREM to wake (Fig. 5B), hippocampal firing decreased significantly particularly among the lowest firing quintiles. These changes resulted in a leftward shift in the firing rate distribution (p = 2.9×10^−18^, Kolmogorov-Smirnov test) and an increase in the CV (ΔCV=0.915 ± 0.137, p = 1.9×10^−7^, Wilcoxon signed-rank test). In the neocortex, on the other hand, higher firing principle neurons increased firing at the transitions from NREM to wake, essentially reversing the change from wake to NREM (Fig. 5A), and producing a significant increase in the CV of the distributions (ΔCV=0.243 ± 0.082, p = 0.019, Wilcoxon signed-rank test). The transitions from REM to wake showed slightly different effects across quintiles (Fig. 5C). In sum, wake and sleep have contrasting effects on the activity of neurons in different quintiles, with sleep states displaying a more homogeneous distribution of firing rates and greater variation among the population during wake.

### Lasting effects of sleep and sleep states on firing rate distributions

These analyses describe a perpetually fluctuating pattern of neuronal activity across sleep and wake transitions, with alternating narrowing and widening of firing rate distributions. We next asked which of these effects persists across longer sleep sequences composed of multiple NREM and REM episodes. First, we analyzed state triplets composed of NREM_i_-REM-NREM_i+1_ or REM_i_-NREM-REM_i+1_ (Fig. 6 A-D) (Grosmark et al., 2012; Miyawaki and Diba, 2016; Watson et al., 2016). Time normalized firing rates and CVs in the triplets further illustrated and confirmed the distribution narrowing and widening effects of NREM and REM epochs, respectively (Fig. 6 A, B), in both brain regions. However, based on these plots it appeared that these distribution changes largely offset and cancelled one another. To better quantify these impressions, we again calculated *DI*s for quintiles in both regions and compared firing rate distributions and CVs. All hippocampal quintiles showed decreased firing between consecutive NREMs interleaved by REM (Fig. 6C; note also that these decreases were more uniform across quintiles than those reported in Miyawaki and Diba (2016) because we have excluded epochs with >20% non-firing cells in the present analyses). Firing rate distributions were slightly but significantly shifted leftward (p =2.7×10^−5^, Kolmogorov-Smirnov test), though CVs were not statistically significant (ΔCV=0.024 ± 0.009, p = 0.066, Wilcoxon signed-rank test). In the frontal cortex, lasting effects of REM on NREM_i+1_ versus NREM_i_ were more subtle and lower in magnitude. Only *DI* of the lowest firing quintile was significantly decreased, and we did not detect differences in the firing rate distributions (p =0.99, Kolmogorov-Smirnov test), or CVs (ΔCV=0.004 ± 0.011, p = 0.52, Wilcoxon signed-rank test). In the REM_i_-NREM-REM_i+1_ triplets, significant changes were not detected in *DI*s, distributions, or CVs from either region.

**Figure 6.**
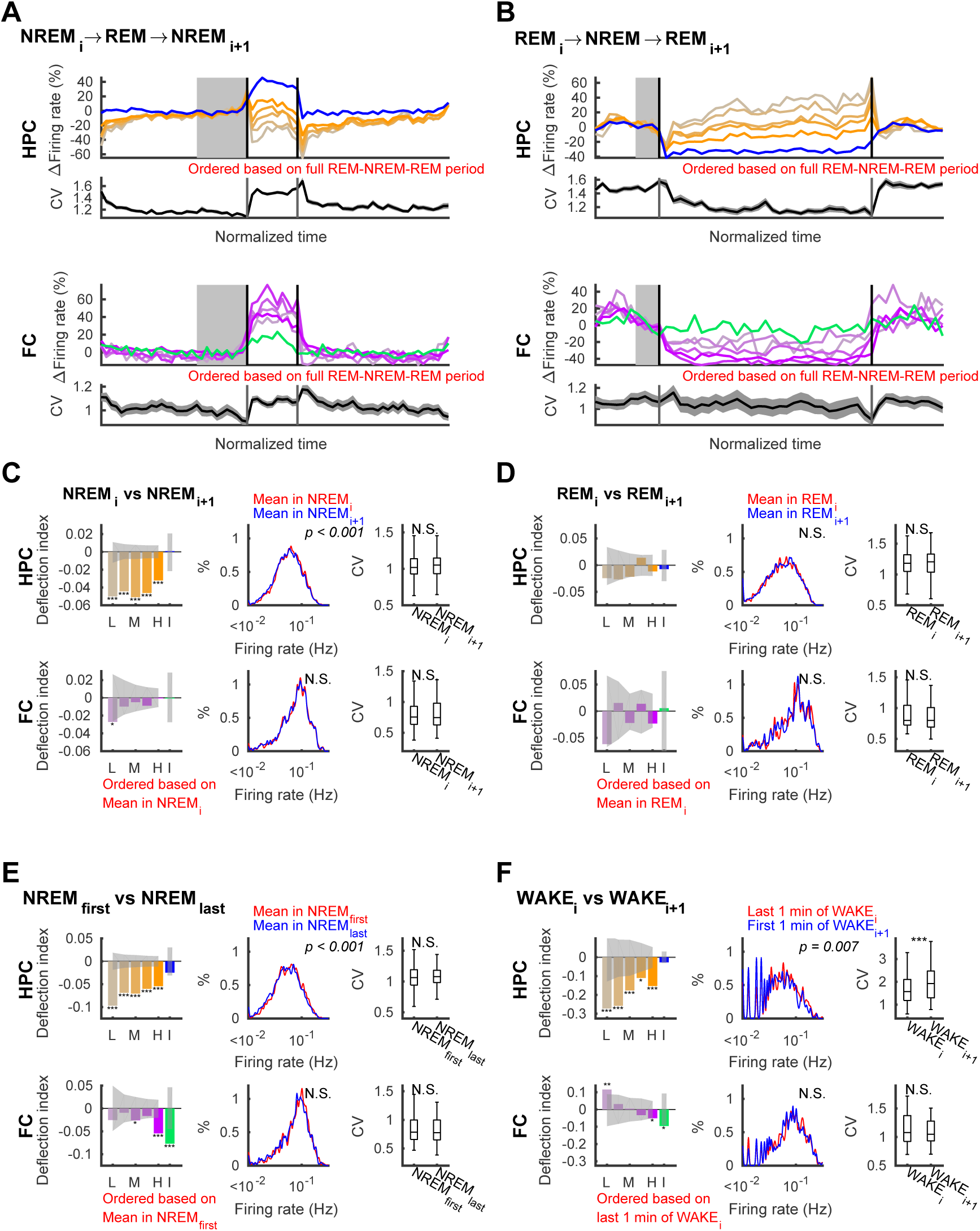
Net effects of sleep of sleep states on neuronal firing distributions – analysis across state triplets. Effect of states as measured by net change from before to after that state. (A, B) Firing rates and coefficient of variation (CV) in the hippocampus (HPC; top panels) and in the frontal cortex (FC; bottom panels) in NREM_i_-REM-NREM_i+1_ triplets (A) and REM_i_-NREM-REM_i+1_ triplets (B) over time normalized for each epoch. Changes in firing rate of each quintile of pyramidal cells (orange shades) and interneurons (blue) in the hippocampus and frontal cortical principal neurons (purple shades) and interneurons (green) are relative to the mean of last third (shown in gray on the top panels and in blue on the bottom panels) of NREM_i_ in (A) and last third of REM_i_ in (B). (C-F) Deflection indices (DI), firing rate distributions, and CV of firing rates in (C) NREMs in NREM-REM-NREM triplets (D), REMs in REM-NREM-REM triplets, between the first and last NREMs in each sleep, and (F) wake periods (last 1-min of WAKE_i_ versus first 1 min of WAKE_i+1_) separated by sleep in the hippocampus (top rows) and in the frontal cortex (bottom rows). L: lowest quintile, M: middle quintile, H: highest quintile, I: interneurons. P-values of Kolmogorov-Smirnov tests are indicated on the middle panels. Changes in CV were tested with the Wilcoxon rank sum test. Error bars and line sheds indicate SEM, sheds on bars indicate 95% confidence intervals of shuffle mean, * p<0.05, ** p<0.01, *** p<0.001, N.S., not significant.

Extending these analyses to first and last NREM in the sleep (separated by longer sequences of alternating REM and NREM), we observed significant decreases across pyramidal cell quintiles in the hippocampus (Fig. 6E). In the frontal cortex, *DI*s were significantly negative only in the middle and highest firing quintiles. These indicate overall firing rate decreases resulting from sleep, consistent with previous reports (Vyazovskiy et al., 2009; Grosmark et al., 2012; Miyawaki and Diba, 2016). Interestingly, the lowest firing quintile decreased most in the hippocampus while the highest firing quintiles decreased most in the frontal cortex. But while firing rate distributions were slightly shifted leftward in the hippocampus (p = 6.1×10^−6^, Kolmogorov-Smirnov test), the difference did not reach significance in the frontal cortex (p = 0.22, Kolmogorov-Smirnov test). Importantly, pairwise comparisons of the CVs did not detect significant changes either in variability in either the hippocampus or the neocortex (ΔCV = 0.039 ± 0.018 (p = 0.093) for hippocampus and −0.010 ± 0.019, (p= 0.731) for frontal cortex, Wilcoxon signed-rank test). These results therefore indicate that distribution changes through multiple sequential REM and NREM states, alternately dispersing and homogenizing firing rates, were counter-balanced throughout sleep in both the hippocampus and the neocortex, despite excitability decreases in both regions in both the population as a whole and in specific quintiles.

Lastly, we compared the last minute of wake before sleep to the first minute of wake following sleep (Fig. 6F). In the hippocampus, *DI*s were significantly negative across quintiles, with lower firing quintiles were more negative than for higher firing quintiles, and firing rate distributions and CVs were significantly different (p = 0.007, Kolmogorov-Smirnov test, and ΔCV= 0.36 ± 0.10, p = 2.3×10^−4^, Wilcoxon signed-rank test). On the other hand, principal neurons in the neocortex showed a significant increase in the lowest firing quintile and a significant decrease in the highest firing quintile, consistent with a narrowed distributed. However, neither firing rate distributions nor CVs were found to be significantly different across sleep (p = 0.97, Kolmogorov-Smirnov test, and ΔCV= −0.044 ± 0.052, p = 0.71, Wilcoxon signed-rank test). These results contradict our expectations based on previous analyses (Watson et al., 2016), which we will address in the Discussion section.

## Discussion

In this work, we aimed to understand how low and high firing hippocampal and frontal cortex neurons are affected during REM and NREM stages of sleep, and upon transitions between these states, to arrive at a better understanding of the function(s) of sleep states in mammals. Since many of our analyses depended on rank-ordering of firing rates and because low-firing and high-firing neurons regress to the mean by chance alone, in this study we designed our analyses to either prevent or correct for this effect. We used two methods; either we based the ordering on the mean firing rates over the entire period being considered, or else we measured all changes relative to a surrogate distribution obtained by random shuffles of the real data. These steps were necessary, because ordering in any selected period produces illusionary normalization in a complementary period and any real changes must be evaluated in contrast to these RTM effects. We found that, in general, sleep states and state transitions do not affect neurons uniformly, but that the changes depend on both the brain region and the relative activity of cells, which likely reflect a combination of neuromodulation of membrane excitability (Graves et al., 2012; Nadim and Bucher, 2014) and sleep-dependent network dynamics involving excitatory and inhibitory synaptic inputs to neurons (Steriade et al., 2001; Timofeev et al., 2001; Dehghani et al., 2016; Niethard et al., 2016; Stringer et al., 2016).

Among the different state dynamics we investigated, NREM sleep was notable in homogenizing excitability across neurons. The transition from REM to NREM produced a greater relative decrease in high-firing cells in both hippocampus and neocortex, with an increase activity of low-firing cells in the hippocampus and a relatively smaller decrease in the neocortex. These changes at the onset of NREM serve to partially homogenize firing across both populations. In the frontal cortex, normalization continued during the NREM episode, and in both regions the coefficient of variation decreased at the onset and further throughout NREM. The onset of NREM was also marked by decreased firing in interneurons in the hippocampus, indicating a shift in the excitation/inhibition balance (see also Stringer et al., 2016). These dynamics across two states characterized by a major shift in cholinergic tone are consistent with the greater relative effect of muscarine on several classes of inhibitory cortical interneurons (Kuchibhotla et al., 2016). Interestingly, atropine, a muscarinic acetylcholine antagonist, also produces increased bursting in hippocampal CA1 pyramidal neurons of lower excitability (“regular spiking”) but decreased bursting in higher excitability (“bursting”) cells (Graves et al., 2012). This suggests that the decreased levels of neuromodulators along with the release from active inhibition allow for a rebalancing of pyramidal cell excitability during NREM sleep.

In contrast, the NREM to REM transition led to greater interneuron spiking and a greater separation of firing between low-firing and high-firing cells, increasing the CV in both regions. These winner-take-all type changes may be implemented in a recurrently connected circuit endowed with inhibition (Yuille and Geiger, 1995; Lee et al., 1999; Rutishauser et al., 2011), such as region CA3, one synapse upstream from our CA1 recordings, or in layer 4 of the neocortex. The shift towards further competition may be supported by increased cholinergic levels during REM sleep that favor feedforward connections, such as from entorhinal cortex to region CA1 (Mizuseki et al., 2011; Schomburg et al., 2014), while neuromodulatory tone in NREM instead favors recurrently-generated activity (Hasselmo, 2006). It is also worth noting that hippocampal interneuron firing patterns across different sleep states closely mirrored those of cortical principal neurons (e.g. see Fig 6A,B), consistent with neocortical control of hippocampal inhibition (Hahn et al., 2006). We also noticed that principal neurons showed relatively dramatic changes (e.g. see Fig. 6A, B) at the transitions between NREM and REM. These transition points may have unique properties: the transitionary period from NREM to REM sleep may in fact be a unique period of “intermediate sleep” that is inundated with both thalamacortical sleep spindles and theta oscillations (Gottesmann, 1992; Watson et al., 2016), while the transitions from REM to NREM are often followed by LOW states and microarousals (Miyawaki et al., 2017).

The net effects of these state transitions, from the first to the last NREM epochs during extended sleep sequences were mostly consistent with our previous reports (Miyawaki and Diba, 2016; Watson et al., 2016), with some notable differences. Here, we find that distribution of the firing rates spread during REM and the homogenization during NREM largely cancel out in both hippocampus and neocortex, yielding a net effect of decreased firing rates in both regions over sleep (Cirelli, 2017). These decreases were seen across all hippocampal quintiles over sleep, but preferentially in lower-firing neurons (Miyawaki and Diba, 2016). In the neocortex, decreases were less pronounced and were specific to high-firing cells, whereas Watson et al. (2016) reported an additional parallel increase in firing of lower-firing neurons. This discrepancy between the present study and Watson et al. (2016) may arise because of two factors: 1) Watson et al. (2016) did not shuffle correct for RTM as in the current *DI* analysis and 2) Watson et al. (2016) did not compare changes across entire episodes of NREM sleep, but rather across bouts or “packets” of NREM. The first and last packets of NREM fall onto the first and last thirds of NREM, and indeed, we observed a narrowing of frontal cortex firing rate distributions within each NREM episode over this period (see Fig. 4A). It should also be noted that the comparison of WAKE_i_ versus WAKE_i+1_ (Fig. 6F) showed simultaneous firing increases in low-firing cells and firing decreases in high-firing cells, consistent with Watson et al. (2016).

We and others have conjectured that the slower firing rate decreases over sleep, on the other hand, are produced by the downscaling of synaptic connections (Vyazovskiy et al., 2009; Grosmark et al., 2012; Tononi and Cirelli, 2014; Miyawaki and Diba, 2016). Network modeling also supports the notion of a strong link between the strength a neuron’s connectivity and its firing rate (Olcese et al., 2010; Lim et al., 2015). Hippocampal changes across sleep (Fig. 6E,F) are consistent with an additive change (e.g. Fig 3C), which indicates hippocampal firing decreased by a similar amount across cells. If the conjecture between synaptic connection and firing rate is correct, the uniform decrease of firing rates could imply a uniform weakening of synaptic connections which effectively improves signal-to-noise in higher-firing cells (Tononi and Cirelli, 2014; see also Cirelli, 2017). A recent study employing scanning electron microscopy of synaptic connections in the cortex supports this analysis; following sleep but not waking, smaller axonal-spine interfaces were observed in the four lower quintiles, with a lesser or no effect in the highest quintile (de Vivo et al., 2017). Higher-firing neurons appear to show the least plasticity, perhaps as a consequence of rigidity or saturated synapses (Grosmark and Buzsaki, 2016). These distinctions may also reflect differences in neuronal subtypes within the CA1 pyramidal layer (Mizuseki et al., 2011; Danielson et al., 2016) that exist throughout the cortex (Molyneaux et al., 2007), though surprisingly in the frontal cortex we saw the greatest decrease in firing across sleep in the higher-firing quintile (but see discussion points below).

Overall, these observations demonstrate a remarkable degree of agreement about the effects of wake and sleep states on neuronal firing in the hippocampus and the frontal neocortex. However, some inter-regional differences were also evident in the responses of quintiles, particularly during the course of NREM episodes and at the transition from NREM to REM. A possible source of differences in hippocampal versus cortical profiles is that the cortex has DOWN states—periods of temporary network silence during NREM—which are not as clearly defined in the hippocampus (Steriade et al., 1993; Isomura et al., 2006). The predominance of DOWN states can potentially account for the relatively decreased firing activity in the neocortex during NREM (see also Vyazovskiy et al., 2009), particularly in the highest firing rate groups as slow waves in NREM develop, and the strong rebound in firing in these quintiles at the onset of REM sleep. On the other hand, LOW states and microarousals at the onset of NREM seem to have stronger suppressive effects on the firing of hippocampal neurons (Miyawaki et al., 2017). These apparent inter-regional differences may also arise because recordings and unit and state detection were performed by different experimenters in different labs. It is worth noting that overall firing rates were higher for the frontal cortex recordings than for the hippocampus recordings, so that lowest quintiles in the frontal cortex fire at similar rates to the middle quintiles in the hippocampus. Hence, sleep states may have effects that depend on absolute rather than relative firing rates and the normalizing and dispersing effects of NREM and REM sleep, respectively, represent broad effects of the neuromodulatory tones under different brain states.

## Acknowledgements

This work was supported by NIH R01MH109170.

## References

Bartram J, Kahn MC, Tuohy S, Paulsen O, Wilson T, Mann EO (2017) Cortical Up states induce the selective weakening of subthreshold synaptic inputs. Nat Commun 8:665. PMCID: PMCPMC5610171 PMID:28939859.

Basu J, Zaremba JD, Cheung SK, Hitti FL, Zemelman BV, Losonczy A, Siegelbaum SA (2016) Gating of hippocampal activity, plasticity, and memory by entorhinal cortex long-range inhibition. Science 351:aaa5694. PMCID: PMCPMC4920085 PMID:26744409.

Brown RE, Basheer R, McKenna JT, Strecker RE, McCarley RW (2012) Control of sleep and wakefulness. Physiol Rev 92:1087–1187. PMCID: PMC3621793 PMID:22811426.

Cheng S, Frank LM (2008) New experiences enhance coordinated neural activity in the hippocampus. Neuron 57:303–313. PMCID: PMC2244590 PMID:18215626.

Cirelli C (2017) Sleep, synaptic homeostasis and neuronal firing rates. Curr Opin Neurobiol 44:72–79. PMCID: PMCPMC5605801 PMID:28399462.

Danielson NB, Zaremba JD, Kaifosh P, Bowler J, Ladow M, Losonczy A (2016) Sublayer-Specific Coding Dynamics during Spatial Navigation and Learning in Hippocampal Area CA1. Neuron 91:652–665. PMCID: PMC4975984 PMID:27397517.

de Vivo L, Bellesi M, Marshall W, Bushong EA, Ellisman MH, Tononi G, Cirelli C (2017) Ultrastructural evidence for synaptic scaling across the wake/sleep cycle. Science 355:507–510. PMCID: PMCPMC5313037 PMID:28154076.

Dehghani N, Peyrache A, Telenczuk B, Le Van Quyen M, Halgren E, Cash SS, Hatsopoulos NG, Destexhe A (2016) Dynamic Balance of Excitation and Inhibition in Human and Monkey Neocortex. Sci Rep 6:23176. PMCID: PMC4793223 PMID:26980663.

Evarts EV (1964) Temporal Patterns of Discharge of Pyramidal Tract Neurons during Sleep and Waking in the Monkey. J Neurophysiol 27:152–171 PMID:14129768.

Gottesmann C (1992) Detection of seven sleep-waking stages in the rat. Neurosci Biobehav Rev 16:31–38 PMID:1553104.

Graves AR, Moore SJ, Bloss EB, Mensh BD, Kath WL, Spruston N (2012) Hippocampal pyramidal neurons comprise two distinct cell types that are countermodulated by metabotropic receptors. Neuron 76:776–789. PMCID: PMC3509417 PMID:23177962.

Grosmark AD, Buzsaki G (2016) Diversity in neural firing dynamics supports both rigid and learned hippocampal sequences. Science 351:1440–1443. PMCID: PMC4919122 PMID:27013730.

Grosmark AD, Mizuseki K, Pastalkova E, Diba K, Buzsaki G (2012) REM sleep reorganizes hippocampal excitability. Neuron 75:1001–1007. PMCID: PMC3608095 PMID:22998869.

Gulati T, Guo L, Ramanathan DS, Bodepudi A, Ganguly K (2017) Neural reactivations during sleep determine network credit assignment. Nat Neurosci 20:1277–1284. PMCID: PMCPMC5808917 PMID:28692062.

Hahn TT, Sakmann B, Mehta MR (2006) Phase-locking of hippocampal interneurons’ membrane potential to neocortical up-down states. Nat Neurosci 9:1359–1361 PMID:17041594.

Hasselmo ME (2006) The role of acetylcholine in learning and memory. Curr Opin Neurobiol 16:710–715. PMCID: PMC2659740 PMID:17011181.

Hengen KB, Lambo ME, Van Hooser SD, Katz DB, Turrigiano GG (2013) Firing rate homeostasis in visual cortex of freely behaving rodents. Neuron 80:335–342. PMCID: PMC3816084 PMID:24139038.

Hengen KB, Torrado Pacheco A, McGregor JN, Van Hooser SD, Turrigiano GG (2016) Neuronal Firing Rate Homeostasis Is Inhibited by Sleep and Promoted by Wake. Cell 165:180–191. PMCID: PMC4809041 PMID:26997481.

Hobson JA, Pace-Schott EF (2002) The cognitive neuroscience of sleep: neuronal systems, consciousness and learning. Nat Rev Neurosci 3:679–693 PMID:12209117.

Isomura Y, Sirota A, Ozen S, Montgomery S, Mizuseki K, Henze DA, Buzsaki G (2006) Integration and segregation of activity in entorhinal-hippocampal subregions by neocortical slow oscillations. Neuron 52:871–882 PMID:17145507.

Koulakov AA, Hromadka T, Zador AM (2009) Correlated connectivity and the distribution of firing rates in the neocortex. J Neurosci 29:3685–3694. PMCID: PMC2784918 PMID:19321765.

Kuchibhotla KV, Gill JV, Lindsay GW, Papadoyannis ES, Field RE, Sten TA, Miller KD, Froemke RC (2016) Parallel processing by cortical inhibition enables context-dependent behavior. Nat Neurosci PMID:27798631.

Lee D, Lin BJ, Lee AK (2012) Hippocampal place fields emerge upon single-cell manipulation of excitability during behavior. Science 337:849–853 PMID:22904011.

Lee DK, Itti L, Koch C, Braun J (1999) Attention activates winner-take-all competition among visual filters. Nat Neurosci 2:375–381 PMID:10204546.

Lim S, McKee JL, Woloszyn L, Amit Y, Freedman DJ, Sheinberg DL, Brunel N (2015) Inferring learning rules from distributions of firing rates in cortical neurons. Nat Neurosci 18:1804–1810. PMCID: PMC4666720 PMID:26523643.

Marder E, Goaillard JM (2006) Variability, compensation and homeostasis in neuron and network function. Nat Rev Neurosci 7:563–574 PMID:16791145.

McCarley RW, Hobson JA (1971) Single neuron activity in cat gigantocellular tegmental field: selectivity of discharge in desynchronized sleep. Science 174:1250–1252 PMID:5133450.

Miyawaki H, Diba K (2016) Regulation of Hippocampal Firing by Network Oscillations during Sleep. Curr Biol 26:893–902. PMCID: PMC4821660 PMID:26972321.

Miyawaki H, Billeh YN, Diba K (2017) Low Activity Microstates During Sleep. Sleep 40 PMID:28431164.

Mizuseki K, Diba K, Pastalkova E, Buzsaki G (2011) Hippocampal CA1 pyramidal cells form functionally distinct sublayers. Nat Neurosci 14:1174–1181. PMCID: PMC3164922 PMID:21822270.

Molyneaux BJ, Arlotta P, Menezes JR, Macklis JD (2007) Neuronal subtype specification in the cerebral cortex. Nat Rev Neurosci 8:427–437 PMID:17514196.

Nadim F, Bucher D (2014) Neuromodulation of neurons and synapses. Curr Opin Neurobiol 29:48–56. PMCID: PMC4252488 PMID:24907657.

Niethard N, Hasegawa M, Itokazu T, Oyanedel CN, Born J, Sato TR (2016) Sleep-Stage-Specific Regulation of Cortical Excitation and Inhibition. Curr Biol 26:2739–2749 PMID:27693142.

Nigam S, Shimono M, Ito S, Yeh FC, Timme N, Myroshnychenko M, Lapish CC, Tosi Z, Hottowy P, Smith WC, Masmanidis SC, Litke AM, Sporns O, Beggs JM (2016) Rich-Club Organization in Effective Connectivity among Cortical Neurons. J Neurosci 36:670–684. PMCID: PMC4719009 PMID:26791200.

Olcese U, Esser SK, Tononi G (2010) Sleep and synaptic renormalization: a computational study. J Neurophysiol 104:3476–3493. PMCID: PMC3007640 PMID:20926617.

Renouard L, Billwiller F, Ogawa K, Clement O, Camargo N, Abdelkarim M, Gay N, Scote-Blachon C, Toure R, Libourel PA, Ravassard P, Salvert D, Peyron C, Claustrat B, Leger L, Salin P, Malleret G, Fort P, Luppi PH (2015) The supramammillary nucleus and the claustrum activate the cortex during REM sleep. Sci Adv 1:e1400177. PMCID: PMCPMC4640625 PMID:26601158.

Roxin A, Brunel N, Hansel D, Mongillo G, van Vreeswijk C (2011) On the distribution of firing rates in networks of cortical neurons. J Neurosci 31:16217–16226 PMID:22072673.

Rutishauser U, Douglas RJ, Slotine JJ (2011) Collective stability of networks of winner-take-all circuits. Neural Comput 23:735–773 PMID:21162667.

Saper CB, Fuller PM, Pedersen NP, Lu J, Scammell TE (2010) Sleep state switching. Neuron 68:1023–1042. PMCID: PMC3026325 PMID:21172606.

Schomburg EW, Fernandez-Ruiz A, Mizuseki K, Berenyi A, Anastassiou CA, Koch C, Buzsaki G (2014) Theta phase segregation of input-specific gamma patterns in entorhinal-hippocampal networks. Neuron 84:470–485. PMCID: PMC4253689 PMID:25263753.

Steriade M, Nunez A, Amzica F (1993) A novel slow (< 1 Hz) oscillation of neocortical neurons in vivo: depolarizing and hyperpolarizing components. J Neurosci 13:3252–3265 PMID:8340806.

Steriade M, Timofeev I, Grenier F (2001) Natural waking and sleep states: a view from inside neocortical neurons. J Neurophysiol 85:1969–1985 PMID:11353014.

Stringer C, Pachitariu M, Okun M, Bartho P, Harris K, Latham P, Sahani M, Lesica N (2016) Inhibitory control of shared variability in cortical networks. BioRxiv oi: http://dx.doi.org/10.1101/041103.

Stuart GJ, Spruston N (2015) Dendritic integration: 60 years of progress. Nat Neurosci 18:1713- 1721 PMID:26605882.

Timofeev I, Grenier F, Steriade M (2001) Disfacilitation and active inhibition in the neocortex during the natural sleep-wake cycle: an intracellular study. Proc Natl Acad Sci U S A 98:1924–1929. PMCID: PMC29358 PMID:11172052.

Tononi G, Cirelli C (2014) Sleep and the price of plasticity: from synaptic and cellular homeostasis to memory consolidation and integration. Neuron 81:12–34 PMID:24411729.

Tsodyks MV, Skaggs WE, Sejnowski TJ, McNaughton BL (1996) Population dynamics and theta rhythm phase precession of hippocampal place cell firing: a spiking neuron model. Hippocampus 6:271–280 PMID:8841826.

Turrigiano G (2011) Too many cooks? Intrinsic and synaptic homeostatic mechanisms in cortical circuit refinement. Annu Rev Neurosci 34:89–103 PMID:21438687.

Vyazovskiy VV, Olcese U, Lazimy YM, Faraguna U, Esser SK, Williams JC, Cirelli C, Tononi G (2009) Cortical firing and sleep homeostasis. Neuron 63:865–878. PMCID: PMC2819325 PMID:19778514.

Watson BO, Levenstein D, Greene JP, Gelinas JN, Buzsaki G (2016) Network Homeostasis and State Dynamics of Neocortical Sleep. Neuron 90:839–852. PMCID: PMC4873379 PMID:27133462.

Weber F, Dan Y (2016) Circuit-based interrogation of sleep control. Nature 538:51–59 PMID:27708309.

Xie X, Hahnloser RHR, Seung HS (2002) Selectively Grouping Neurons in Recurrent Networks of Lateral Inhibition. Neural Comput 14:2627–2646.

Yassin L, Benedetti BL, Jouhanneau JS, Wen JA, Poulet JF, Barth AL (2010) An embedded subnetwork of highly active neurons in the neocortex. Neuron 68:1043–1050. PMCID: PMC3022325 PMID:21172607.

Yuille AL, Geiger D (1995) Winner-take-all mechanisms. In: Handbook of Brain Theory and Neural Networks (Arbib MA, ed), pp 1–1056: Mit Press.

